# Enhanced proteome profiling of human cerebrospinal fluid using a commercial plasma enrichment strategy

**DOI:** 10.1101/2024.10.07.616086

**Authors:** Eva Borràs, Federica Anastasi, Olga Pastor, Marc Suárez-Calvet, Eduard Sabidó

## Abstract

Cerebrospinal fluid (CSF) is a valuable liquid biopsy for identifying protein biomarkers in neurological diseases, yet its proteome profiling faces challenges due to the large dynamic range of protein abundances. In this study, we assessed the effectiveness of a commercial enrichment strategy, initially developed for plasma samples, in enhancing the detection of low-abundance proteins in human CSF. We demonstrate significant improvements in protein identification and coverage depth while maintaining high reproducibility and low coefficients of variation. These findings underscore the potential of this enrichment strategy to facilitate rapid and sensitive CSF analysis, advancing biomarker discovery in neurological research.

---

Cerebrospinal fluid (CSF) is a valuable liquid biopsy for neurological diseases research to identify protein biomarkers for diagnostic purposes,^1^ predict treatment response,^2^ and to understand the molecular mechanisms involved in disease progression.^3–7^ However, profiling the human CSF proteome by mass spectrometry faces two main analytical challenges. Firstly, the broad dynamic range of protein abundances commonly seen in liquid biopsies, which hinders the sensitivity and the ability to measure low-abundant proteins. Secondly, the need to prepare and analyze hundreds of human samples in a reproducible and timely manner. Indeed, there has been a trade-off among sensitivity, number of detected analytes, number of samples, sample amount, and data quality. Multiplexed immunoassays and aptamer-based approaches have recently been developed to address these analytical challenges;^8,9^ however, they are limited by their targeted nature, as they can only identify a pre-selected panel of proteins. Recent advancements in mass spectrometry instrumentation have facilitated the rapid acquisition of liquid biopsies,^10–13^ which is essential for analyzing large patient cohorts in translational clinical projects. Concurrently, several sample preparation commercial solutions have been introduced to address the challenges associated with the extensive dynamic range of liquid biopsies, and enhance in-depth protein identification and increase sample throughput. These new approaches offer efficient alternatives to existing classical strategies like antibody-based protein depletion and sample fractionation strategies,^14,15^ and include the use of nanoparticle protein coronas,^16,17^ hyper-porous strong-anion exchange magnetic microparticles,^18^ and seeded precipitation on paramagnetic beads to enrich low abundant proteins.^19^ Despite their demonstrated superior performance in plasma samples,^20^ their potential to enhance the analytical sensitivity of CSF remains to be established. In this study, we assessed the PreOmics ENRICH-iST kit protocol—originally designed to reduce the dynamic range in plasma samples—for processing CSF, and compared its performance to the analysis of neat CSF samples. We assessed various amounts of CSF starting material, and evaluated the sensitivity and reproducibility of this enrichment strategy.

Initially, a pool of human CSF samples was obtained from two individuals with Alzheimer’s disease dementia and mild cognitive impairment, and it was consistently used in all subsequent experiments. The pooled CSF sample was processed in varying amounts, employing a tryptic digestion directly on the neat CSF samples, or after an enrichment procedure using paramagnetic beads. For the analysis of neat CSF, the PreOmics iST-BCT 8x kit was used, processing 10 μL of CSF according to the manufacturer’s protocol to obtain the peptide mix prior to analysis by liquid chromatography coupled to mass spectrometry (LC-MS). For the enrichment of low-abundant proteins, we employed the ENRICH iST-BCT 8x kit, which had been previously optimized for 20 μL of plasma.^19^ Importantly, CSF samples have a protein concentration approximately 100 times lower than that of neat plasma (plasma: ∼50 μg/μL; CSF: ∼0,25-0.5 μg/μL), thus requiring the evaluation of higher volumes of CSF as part of our optimization process. We processed three different starting amounts of CSF representing a volume increase ranging from 2.5-to 25-fold compared to the one recommended for plasma (**Figure 1A**). The binding buffer used during the enrichment process had also to be adapted. We therefore assessed several amounts of binding buffer in conjunction with different volumes of CSF. The specific conditions tested included 50 μL of CSF in 70 μL of binding buffer, 150 μL of CSF in 200 μL of binding buffer, and 500 μL of CSF in 700 μL of binding buffer. The volumes tested were carefully chosen to maximize protein processing while minimizing sample usage due to the intrinsic limited availability of human CSF samples, and the need to ensure that the selected volumes fit within the volume constraints of the kit. In all cases, 10% of the digested enriched eluate was loaded into the analytical column. For the neat CSF, *ca*. 1 μg was loaded into the column in the Orbitrap Eclipse system, and *ca*. 100 ng were loaded into the column in the Orbitrap Astral, following the recommended amounts for each platform. All experiments were performed in technical triplicate starting from the same pool of human CSF sample.

**Figure 1:**
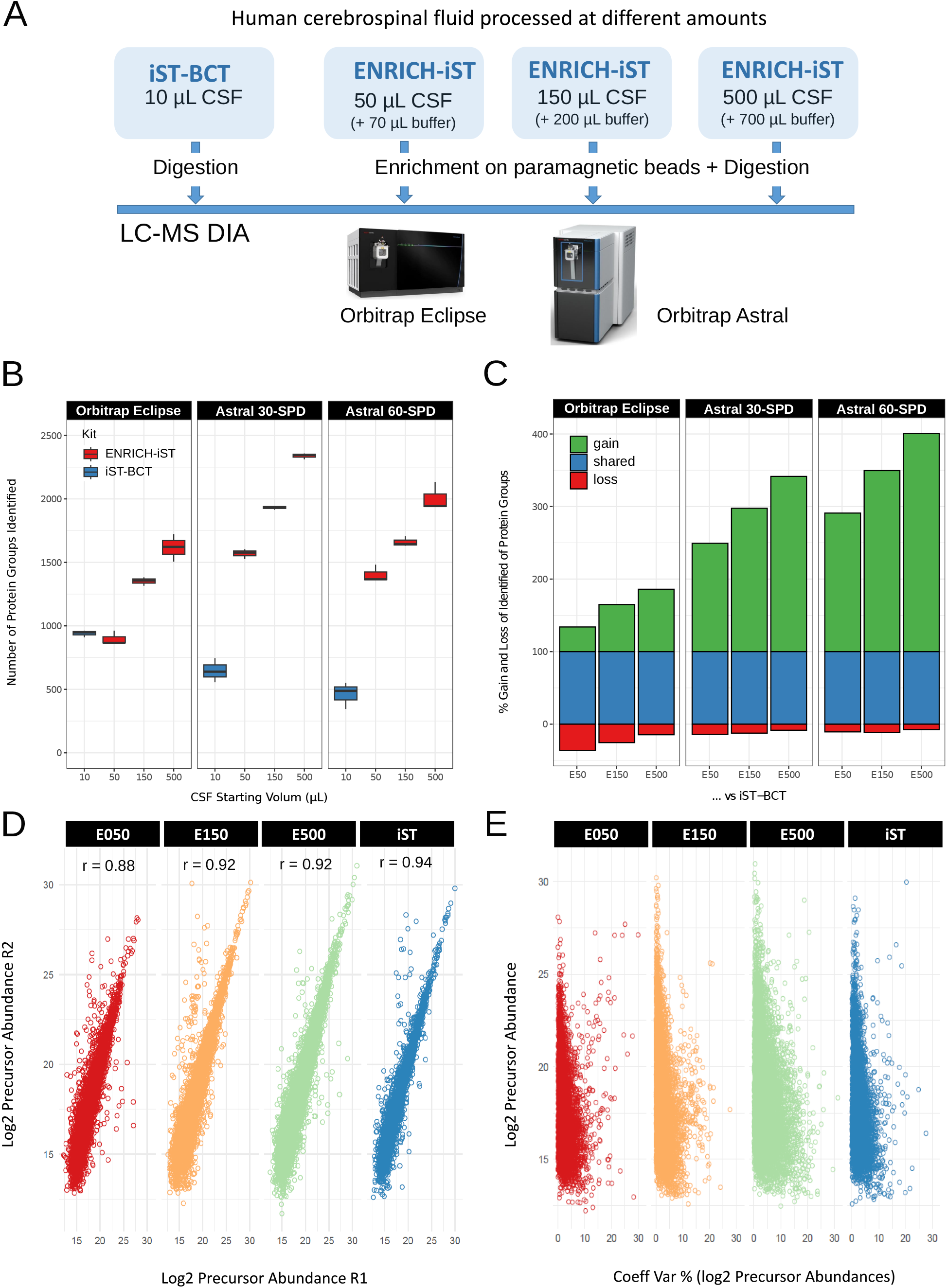
A) Overall view of the experiments performed with cerebrospinal fluid (CSF) in this work; B) Number of protein groups identified in neat and enriched CSF samples in the different mass spectrometry platforms; C) Percentage of gain and loss protein groups identification in each of the enriched samples compared to the neat CSF sample (iST-BCT). Different starting volumes for the enrichment protocol were tested: 50 μL (E50), 150 μL (E150), and 500 μL (E500); D) Correlation of precursor abundances (logarithmic scale) among technical replicates in different enriched CSF volumes and neat CSF (iST-BCT); and E) Coefficient of variation for precursor abundance among triplicate measurements in different enriched CSF volumes and neat CSF (iST-BCT).

Digested and enriched samples were analyzed by LC-MS using data-independent acquisition across two different instruments: an Orbitrap Eclipse Tribrid, and an Orbitrap Astral. Chromatography settings and data-acquisition schema were tailored for each platform (**Supporting Information**). Acquired spectra were analyzed using a library-free strategy with DIA-NN (Neural networks and interference correction enable deep proteome coverage in high throughput) (v1.8.1). The data were searched against a Swiss-Prot human database (as in April 2023) plus a list of common contaminants, and all the corresponding decoy entries. For peptide identification trypsin was chosen as enzyme and up to one miscleavage was allowed. Oxidation of methionine was used as variable modification whereas carbamidomethylation on cysteines was set as a fixed modification. False discovery rate (FDR) was set to a maximum of 1% at peptide and protein level. Precursor and fragment ion m/z mass range were adjusted to 500-900 and 350-1850, respectively. For peptide quantification match-between-runs was enabled, protein inference was set to ‘Protein names (from FASTA)’ with ‘Heuristic protein inference’ option and the quantification strategy was set to ‘Robust LC (high precision)’. Default settings were used for the other parameters. The mass spectrometry proteomics data have been deposited to the ProteomeXchange Consortium via the PRIDE partner repository with the dataset identifier PXD055853.^21^

Results were initially analyzed in terms of protein identifications to determine whether the enrichment procedure led to the identification of more analytes compared to the neat CSF analysis (**Figure 1B** and **1C, Supplementary Table S1**). In the results generated from the Orbitrap Eclipse, we observed an increase in protein identification in the enriched samples when using at least 150 μL and 500 μL of starting CSF material. Similarly, in the Orbitrap Astral, there was a significant increase in the number of protein identifications when enriching CSF samples no matter the starting volume (50 μL, 150 μL, 500 μL), nor the data acquisition strategy used (30 SPD, 60 SPD). In all cases, the number of identified protein groups in enriched samples increased with the initial volume of CSF increased, indicating that the enrichment procedure had not reached saturation. However, larger CSF volumes were not evaluated due to limitations in sample availability, constraints of the kit, and, importantly, because using larger volumes would not provide clinically translatable results. In the case of the Orbitrap Eclipse, no significant increase in the number of identified proteins was observed when enriching low amounts of CSF (i.e., 50 μL) compared to the neat CSF. In contrast, a clear difference was noted in the Orbitrap Astral between neat CSF and low amounts of CSF (i.e., 50 μL). This observation is likely due to the excellent analytical performance of the Orbitrap Eclipse system when operated with high amounts of neat CSF in combination with the use of a 50-cm column and a 2-hour gradient. Conversely, in the Orbitrap Astral, the lower loading amounts and shorter gradients used (**Supporting Information**) probably limited the comprehensive analysis of neat CSF. In these high-throughput analyses, the use of the enrichment process results in a considerable gain in the number of identified proteins, even when starting the enrichment procedure with only 50 μL of CSF. In the most favorable conditions tested, up to 2,616 protein groups and 23,875 peptide precursors were identified when analyzing all 3 replicates together (30 SPD, Orbitrap Astral, 500 μL of CSF). Nevertheless, utilizing 150 μl of CSF in conjunction with short data acquisition gradients (e.g., 30 SPD) probably represents the approach with the highest translational potential, as it effectively balances analysis time, sample volume, and the number of identified peptide precursors and protein groups. Interestingly, when focusing on specific proteins previously linked to diseases such as multiple sclerosis, amyotrophic lateral sclerosis, Alzheimer’s, Parkinson’s disease, and other neurodegenerative diseases, we observed a preferential identification of these proteins in samples that had been enriched with paramagnetic beads (**Table 1**). It is important to note that some of the proteins of interest are typically present only in the late stages of certain diseases (e.g. Frataxin), which explains their absence in the CSF samples from the donors used in this study, regardless of the strategy employed.

Beyond the identification of proteins and peptides, it is essential that the enrichment procedure is reproducible to ensure consistent analysis of the CSF. To this end, we assessed the reproducibility of the CSF enrichment procedure by analyzing technical replicates and calculating the coefficients of variation for precursor abundances. We found a strong linear correlation among replicates, with an r value between 0.88-0.92 for the enriched samples analyzed on the Orbitrap Eclipse, comparable to the correlation observed with neat CSF samples with r values of *ca*. 0.94 (**Figure 1D**). Additionally, the coefficients of variation were predominantly below 15% across all conditions tested, and although a slight increase in variation was noted with higher volumes of enriched CSF, the results remained consistent with those obtained from neat CSF samples (**Figure 1E**).

In conclusion, our study successfully evaluated the use of the PreOmics ENRICH-iST kit, an enrichment sample preparation strategy, originally designed for plasma, for processing CSF samples. By adapting this commercially available enrichment strategy, we significantly enhanced proteome coverage and depth in human CSF while ensuring high technical reproducibility and low coefficients of variation. A known limitation of this study is the inability to conduct a comprehensive assessment of additional analytical conditions that would be of interest from a strictly analytical perspective. This constraint arises from the limited availability of samples and the ethical considerations associated with the use of human CSF. However, our findings underscore the potential of such enrichment methods to improve the identification of low-abundant proteins in human CSF, which is crucial for advancing biomarker discovery and clinical applications in neurological disease research. The integration of optimized sample preparation techniques with cutting-edge mass spectrometry instrumentation will certainly facilitate rapid and sensitive analysis of CSF samples, thereby supporting large-scale studies aimed at understanding complex neurological conditions.

## Acknowledgements

The authors would like to express their most sincere gratitude to the staff of the Neurology department at Hospital del Mar and the BIODEGMAR participants and relatives without whom this research would have not been possible. We acknowledge support of the Spanish Ministry of Science and Innovation through the Centro de Excelencia Severo Ochoa (CEX2020-001049-S grant funded by MCIN/AEI/10.13039/501100011033) and PID2020-115092GB-I00 funded by AEI/10.13039/501100011033, and the Generalitat de Catalunya through the CERCA programme and the Departament de Recerca i Universitats (2021-SGR2021-01225). This result is part of a project that has received funding from the European Union’s Horizon 2020 research and innovation programme under the Marie Sklodowska-Curie grant agreement No 956148. The CRG/UPF Proteomics Unit is part of the Spanish Infrastructure for Omics Technologies (ICTS OmicsTech). FA receives funding from the JDC2022-049347-I grant, funded by the MCIU/AEI/10.13039/501100011033 and the European Union NextGenerationEU/PRTR. MSC receives funding from the European Research Council (ERC) under the European Union’s Horizon 2020 research and innovation program (Grant agreement No. 948677), the Instituto de Salud Carlos III through the projects PI19/00155 and PI22/00456 (Co-funded by European Regional Development Fund (FEDER) “A way to make Europe”), and receives the support of a fellowship from “la Caixa” Foundation (ID 100010434) and from the European Union’s Horizon 2020 research and innovation programme under the Marie Skłodowska-Curie grant agreement No 847648 (fellowship code LCF/BQ/PR21/11840004).

## Ethical Statement

The BIODEGMAR study was approved by the Independent Ethics Committee “Parc de Salut Mar”, Barcelona (CEIC PSMAR, project code 2018/7805I). All participants from BIODEGMAR provided informed consent.

## Notes

The authors declare no competing financial interest. Note that the ENRICH-iST kits used in this work were provided free-of-charge by PreOmics Gmbh.

## Tables

**Table 1:** Detection of specific biomarkers and relevant proteins associated with neurological diseases in both neat and enriched cerebrospinal fluid.

## Supplementary Information

**Materials and Methods:** Detailed description of the LC-MSMS data acquisition methods.

**Supplementary Table S1:** List of peptide precursors and protein groups identified in each condition tested.

## Bibliography

(1) Cantó, E.; Tintoré, M.; Villar, L. M.; Borrás, E.; Alvarez-Cermeño, J. C.; Chiva, C.; Sabidó, E.; Rovira, A.; Montalban, X.; Comabella, M. Validation of Semaphorin 7A and Ala-β-His-Dipeptidase as Biomarkers Associated with the Conversion from Clinically Isolated Syndrome to Multiple Sclerosis. J Neuroinflammation 2014, 11, 181. 10.1186/s12974-014-0181-8.

(2) Faravelli, I.; Gagliardi, D.; Abati, E.; Meneri, M.; Ongaro, J.; Magri, F.; Parente, V.; Petrozzi, L.; Ricci, G.; Farè, F.; Garrone, G.; Fontana, M.; Caruso, D.; Siciliano, G.; Comi, G. P.; Govoni, A.; Corti, S.; Ottoboni, L. Multi-Omics Profiling of CSF from Spinal Muscular Atrophy Type 3 Patients after Nusinersen Treatment: A 2-Year Follow-up Multicenter Retrospective Study. Cell Mol Life Sci 2023, 80 (8), 241. 10.1007/s00018-023-04885-7.

(3) Fissolo, N.; Matute-Blanch, C.; Osman, M.; Costa, C.; Pinteac, R.; Miró, B.; Sanchez, A.; Brito, V.; Dujmovic, I.; Voortman, M.; Khalil, M.; Borràs, E.; Sabidó, E.; Issazadeh-Navikas, S.; Montalban, X.; Comabella Lopez, M. CSF SERPINA3 Levels Are Elevated in Patients With Progressive MS. Neurol Neuroimmunol Neuroinflamm 2021, 8 (2), e941. 10.1212/NXI.0000000000000941.

(4) Comabella, M.; Sastre-Garriga, J.; Borras, E.; Villar, L. M.; Saiz, A.; Martínez-Yélamos, S.; García-Merino, J. A.; Pinteac, R.; Fissolo, N.; Sánchez López, A. J.; Costa-Frossard, L.; Blanco, Y.; Llufriu, S.; Vidal-Jordana, A.; Sabidó, E.; Montalban, X. CSF Chitinase 3-Like 2 Is Associated With Long-Term Disability Progression in Patients With Progressive Multiple Sclerosis. Neurol Neuroimmunol Neuroinflamm 2021, 8 (6), e1082. 10.1212/NXI.0000000000001082.

(5) Lleó, A.; Núñez-Llaves, R.; Alcolea, D.; Chiva, C.; Balateu-Paños, D.; Colom-Cadena, M.; Gomez-Giro, G.; Muñoz, L.; Querol-Vilaseca, M.; Pegueroles, J.; Rami, L.; Lladó, A.; Molinuevo, J. L.; Tainta, M.; Clarimón, J.; Spires-Jones, T.; Blesa, R.; Fortea, J.; Martínez-Lage, P.; Sánchez-Valle, R.; Sabidó, E.; Bayés, À.; Belbin, O. Changes in Synaptic Proteins Precede Neurodegeneration Markers in Preclinical Alzheimer’s Disease Cerebrospinal Fluid. Mol Cell Proteomics 2019, 18 (3), 546–560. 10.1074/mcp.RA118.001290.

(6) Tristán-Noguero, A.; Borràs, E.; Molero-Luis, M.; Wassenberg, T.; Peters, T.; Verbeek, M. M.; Willemsen, M.; Opladen, T.; Jeltsch, K.; Pons, R.; Thony, B.; Horvath, G.; Yapici, Z.; Friedman, J.; Hyland, K.; Agosta, G. E.; López-Laso, E.; Artuch, R.; Sabidó, E.; García-Cazorla, À. Novel Protein Biomarkers of Monoamine Metabolism Defects Correlate with Disease Severity. Mov Disord 2020. 10.1002/mds.28362.

(7) Tijms, B. M.; Vromen, E. M.; Mjaavatten, O.; Holstege, H.; Reus, L. M.; van der Lee, S.; Wesenhagen, K. E. J.; Lorenzini, L.; Vermunt, L.; Venkatraghavan, V.; Tesi, N.; Tomassen, J.; den Braber, A.; Goossens, J.; Vanmechelen, E.; Barkhof, F.; Pijnenburg, Y. A. L.; van der Flier, W. M.; Teunissen, C. E.; Berven, F. S.; Visser, P. J. Cerebrospinal Fluid Proteomics in Patients with Alzheimer’s Disease Reveals Five Molecular Subtypes with Distinct Genetic Risk Profiles. Nat Aging 2024, 4 (1), 33–47. 10.1038/s43587-023-00550-7.

(8) Ellington, A. D.; Szostak, J. W. In Vitro Selection of RNA Molecules That Bind Specific Ligands. Nature 1990, 346 (6287), 818–822. 10.1038/346818a0.

(9) Fredriksson, S.; Gullberg, M.; Jarvius, J.; Olsson, C.; Pietras, K.; Gústafsdóttir, S. M.; Ostman, A.; Landegren, U. Protein Detection Using Proximity-Dependent DNA Ligation Assays. Nat Biotechnol 2002, 20 (5), 473–477. 10.1038/nbt0502-473.

(10) Stewart, H. I.; Grinfeld, D.; Giannakopulos, A.; Petzoldt, J.; Shanley, T.; Garland, M.; Denisov, E.; Peterson, A. C.; Damoc, E.; Zeller, M.; Arrey, T. N.; Pashkova, A.; Renuse, S.; Hakimi, A.; Kühn, A.; Biel, M.; Kreutzmann, A.; Hagedorn, B.; Colonius, I.; Schütz, A.; Stefes, A.; Dwivedi, A.; Mourad, D.; Hoek, M.; Reitemeier, B.; Cochems, P.; Kholomeev, A.; Ostermann, R.; Quiring, G.; Ochmann, M.; Möhring, S.; Wagner, A.; Petker, A.; Kanngiesser, S.; Wiedemeyer, M.; Balschun, W.; Hermanson, D.; Zabrouskov, V.; Makarov, A. A.; Hock, C. Parallelized Acquisition of Orbitrap and Astral Analyzers Enables High-Throughput Quantitative Analysis. Anal Chem 2023, 95 (42), 15656–15664. 10.1021/acs.analchem.3c02856.

(11) Heil, L. R.; Damoc, E.; Arrey, T. N.; Pashkova, A.; Denisov, E.; Petzoldt, J.; Peterson, A. C.; Hsu, C.; Searle, B. C.; Shulman, N.; Riffle, M.; Connolly, B.; MacLean, B. X.; Remes, P. M.; Senko, M. W.; Stewart, H. I.; Hock, C.; Makarov, A. A.; Hermanson, D.; Zabrouskov, V.; Wu, C. C.; MacCoss, M. J. Evaluating the Performance of the Astral Mass Analyzer for Quantitative Proteomics Using Data-Independent Acquisition. J Proteome Res 2023, 22 (10), 3290–3300. 10.1021/acs.jproteome.3c00357.

(12) Mun, D.-G.; Budhraja, R.; Bhat, F. A.; Zenka, R. M.; Johnson, K. L.; Moghekar, A.; Pandey, A. Four-Dimensional Proteomics Analysis of Human Cerebrospinal Fluid with Trapped Ion Mobility Spectrometry Using PASEF. Proteomics 2023, 23 (10), e2200507. 10.1002/pmic.202200507.

(13) Vitko, D.; Chou, W.-F.; Nouri Golmaei, S.; Lee, J.-Y.; Belthangady, C.; Blume, J.; Chan, J. K.; Flores-Campuzano, G.; Hu, Y.; Liu, M.; Marispini, M. A.; Mora, M. G.; Ramaswamy, S.; Ranjan, P.; Williams, P. B.; Zawada, R. J. X.; Ma, P.; Wilcox, B. E. timsTOF HT Improves Protein Identification and Quantitative Reproducibility for Deep Unbiased Plasma Protein Biomarker Discovery. J Proteome Res 2024, 23 (3), 929–938. 10.1021/acs.jproteome.3c00646.

(14) Lee, P. Y.; Osman, J.; Low, T. Y.; Jamal, R. Plasma/Serum Proteomics: Depletion Strategies for Reducing High-Abundance Proteins for Biomarker Discovery. Bioanalysis 2019, 11 (19), 1799–1812. 10.4155/bio-2019-0145.

(15) Tam, S. W.; Pirro, J.; Hinerfeld, D. Depletion and Fractionation Technologies in Plasma Proteomic Analysis. Expert Rev Proteomics 2004, 1 (4), 411–420. 10.1586/14789450.1.4.411.

(16) Blume, J. E.; Manning, W. C.; Troiano, G.; Hornburg, D.; Figa, M.; Hesterberg, L.; Platt, T. L.; Zhao, X.; Cuaresma, R. A.; Everley, P. A.; Ko, M.; Liou, H.; Mahoney, M.; Ferdosi, S.; Elgierari, E. M.; Stolarczyk, C.; Tangeysh, B.; Xia, H.; Benz, R.; Siddiqui, A.; Carr, S. A.; Ma, P.; Langer, R.; Farias, V.; Farokhzad, O. C. Rapid, Deep and Precise Profiling of the Plasma Proteome with Multi-Nanoparticle Protein Corona. Nat Commun 2020, 11 (1), 3662. 10.1038/s41467-020-17033-7.

(17) Ashkarran, A. A.; Gharibi, H.; Voke, E.; Landry, M. P.; Saei, A. A.; Mahmoudi, M. Measurements of Heterogeneity in Proteomics Analysis of the Nanoparticle Protein Corona across Core Facilities. Nat Commun 2022, 13 (1), 6610. 10.1038/s41467-022-34438-8.

(18) Wu, C. C.; Tsantilas, K. A.; Park, J.; Plubell, D.; Sanders, J. A.; Naicker, P.; Govender, I.; Buthelezi, S.; Stoychev, S.; Jordaan, J.; Merrihew, G.; Huang, E.; Parker, E. D.; Riffle, M.; Hoofnagle, A. N.; Noble, W. S.; Poston, K. L.; Montine, T. J.; MacCoss, M. J. Mag-Net: Rapid Enrichment of Membrane-Bound Particles Enables High Coverage Quantitative Analysis of the Plasma Proteome. bioRxiv 2024, 2023.06.10.544439. 10.1101/2023.06.10.544439.

(19) Resources | Preomics. https://www.preomics.com/resources (accessed 2024-08-30).

(20) Soni, R. K. Frontiers in Plasma Proteome Profiling Platforms: Innovations and Applications. Clin Proteomics 2024, 21 (1), 43. 10.1186/s12014-024-09497-2.

(21) Perez-Riverol, Y.; Bai, J.; Bandla, C.; García-Seisdedos, D.; Hewapathirana, S.; Kamatchinathan, S.; Kundu, D. J.; Prakash, A.; Frericks-Zipper, A.; Eisenacher, M.; Walzer, M.; Wang, S.; Brazma, A.; Vizcaíno, J. A. The PRIDE Database Resources in 2022: A Hub for Mass Spectrometry-Based Proteomics Evidences. Nucleic Acids Res 2022, 50 (D1), D543–D552. 10.1093/nar/gkab1038.

