## Supporting Information for "Enhanced proteome profiling of human cerebrospinal fluid using a commercial plasma enrichment strategy"

Technical Note

### Methods

*Chromatographic and mass spectrometric analysis*

**Orbitrap Eclipse**

Samples were analyzed in an Orbitrap Eclipse mass spectrometer (Thermo Fisher Scientific) coupled to an EASY-nLC 1200 (Thermo Fisher Scientific). Peptides were loaded directly onto the analytical column (50 cm length, 75 μm inner diameter, 2 μm C18 particles, Thermo Fisher Scientific) and were separated by reversed-phase chromatography with a 300 nL/min flow and a 120-minute gradient. The instrument was operated in data-independent acquisition mode, with a full MS scans over a mass range of m/z 500-900 with detection in the Orbitrap at a resolution of 120,000. The auto gain control (AGC) was set to 1e6 and a maximum injection time of 246ms was used. In each cycle of data-independent acquisition analysis, 40 windows of 10 Da each were used to isolate and fragment all precursor ions with normalized higher-energy collisional dissociation (HCD) of 28%. MS2 scan range was set from 350 to 1850 m/z with detection in the Orbitrap at a resolution of 30,000. The AGC was set to 1E6 and a maximum injection time of 54 ms was used.

**Orbitrap Astral (Method 30-SPD)**

Samples were analyzed in an Orbitrap Astral mass spectrometer (Thermo Fisher Scientific) coupled to a Vanquish Neo UHPLC (Thermo Fisher Scientific). Peptides were loaded directly onto the analytical column (25 cm length, 75 μm inner diameter, 1.7 μm C18 particles, Aurora Ultimate TS, Ionopticks) and were separated by reversed-phase chromatography with a 500nL/min flow and a 33.3-minute gradient. The instrument was operated in data-independent acquisition mode, with a full MS scans over a mass range of m/z 380-980 with detection in the Orbitrap at a resolution of 240,000. The auto gain control (AGC) was set to 500% and a maximum injection time of 5ms was used. In each cycle of data-independent acquisition analysis, 100 windows of 6 Da each were used to isolate and fragment all precursor ions with normalized higher-energy collisional dissociation (HCD) of 25%. MS2 scan range was set from 150 to 2000 m/z with detection in the Astral. The AGC was set to 500% and a maximum injection time of 10 ms was used.

**Orbitrap Astral (Method 60-SPD)**

Samples were analyzed in an Orbitrap Astral mass spectrometer (Thermo Fisher Scientific) coupled to a Vanquish Neo UHPLC (Thermo Fisher Scientific). Peptides were firstly loaded into a 300um ID x 5mm C18 PepMap Neo trap column and then eluted onto the analytical column (15 cm length, 150 μm inner diameter, 2 μm C18 particles, Easy-Spray, Thermo Fisher Scientific) and were separated by reversed-phase chromatography with a 800nL/min flow and a 20.9 minute gradient. The instrument was operated in data-independent acquisition mode, with a full MS scans over a mass range of m/z 380-980 with detection in the Orbitrap at a resolution of 240,000. The auto gain control (AGC) was set to 500% and a maximum injection time of 5ms was used. In each cycle of data-independent acquisition analysis, 200 windows of 3 Da each were used to isolate and fragment all precursor ions with normalized higher-energy collisional dissociation (HCD) of 25%. MS2 scan range was set from 150 to 2000 m/z with detection in the Astral. The AGC was set to 500% and a maximum injection time of 7 ms was used.
